## Supplementary material for "Structural insights into DNA Topoisomerase II of African Swine Fever Virus": Supplementary.pdf

J. Cong *et al.*

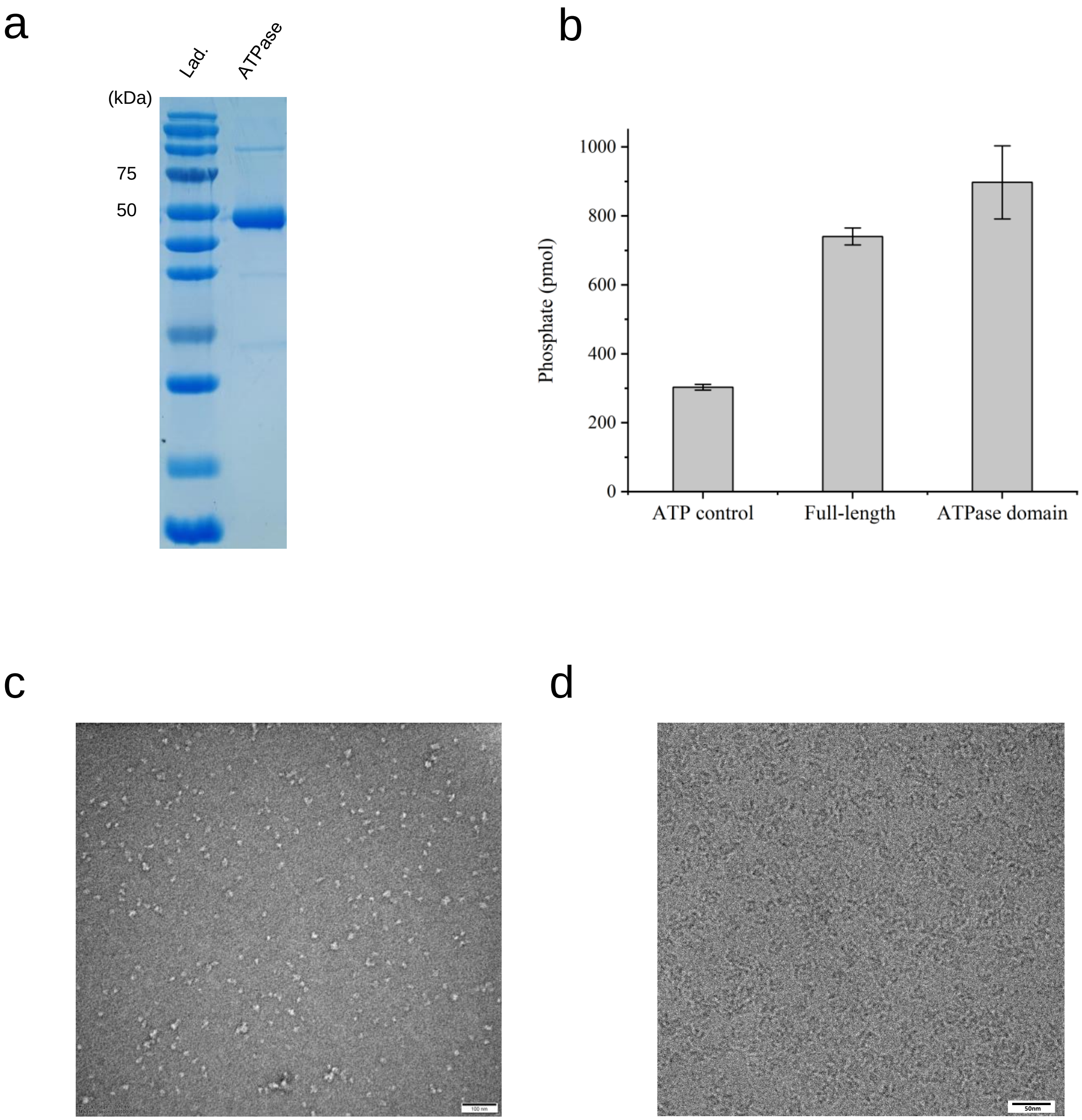

**Supplementary Figure 1: Biochemical assays and negative staining/cryo-EM micrograph. a** SDS-PAGE analysis of the purified truncated ATPase domain. **b** In vitro ATPase activity of the purified full-length pP1192R and truncated ATPase domain. The y-axis represents the total amount of free phosphate ions measured in the solution over 30 minutes. **c** A representative negative staining micrograph. **d** A representative cryo-EM micrograph.

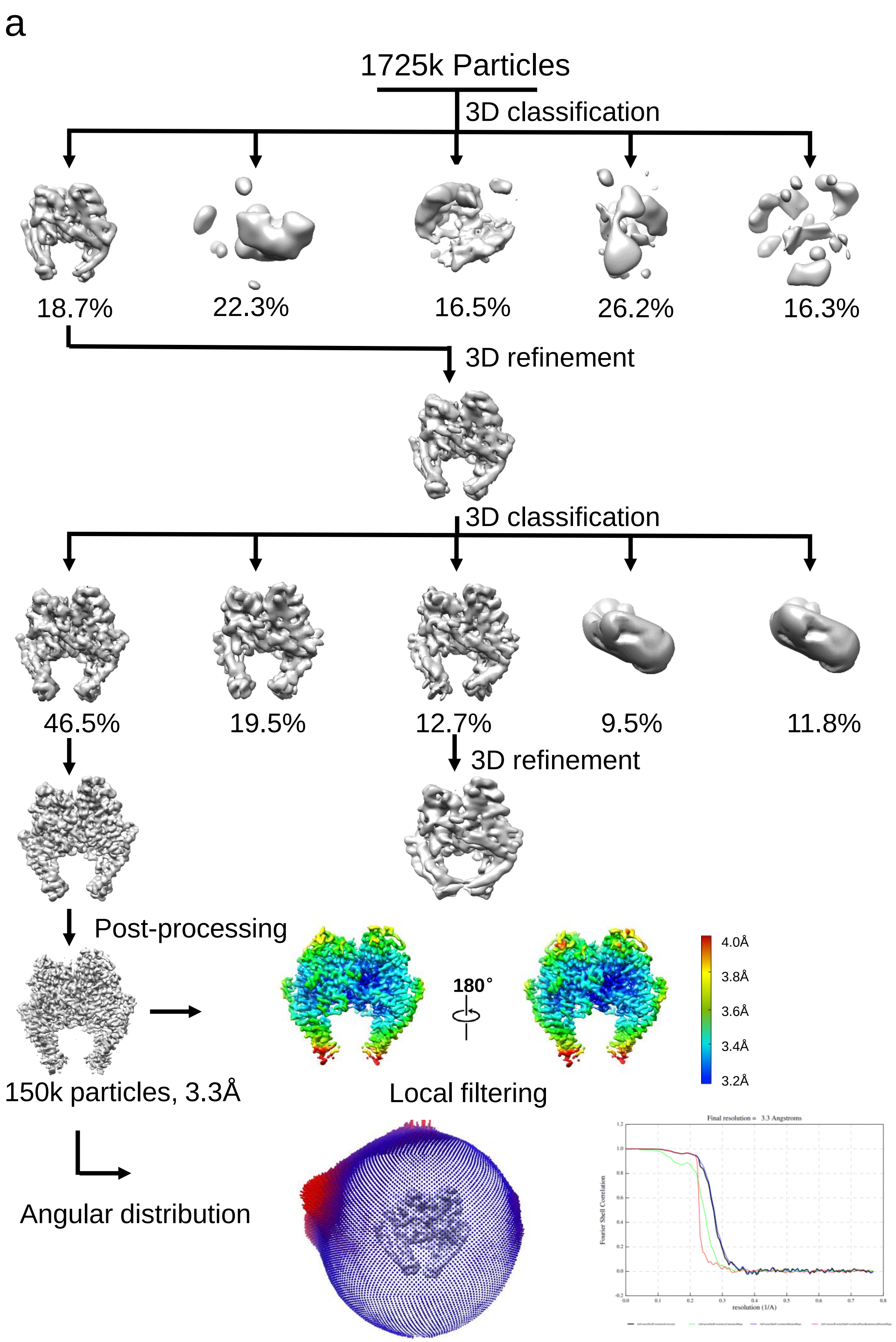
a
1725k Particles
3D classification
18.7%
22.3%
16.5%
26.2%
16.3%
3D refinement
3D classification
46.5%
19.5%
12.7%
9.5%
11.8%
3D refinement
Post-processing
150k particles, 3.3Å
180°
4.0Å
3.8Å
3.6Å
3.4Å
3.2Å
Local filtering
Angular distribution
Final resolution = 3.3 Angstroms
Fourier Shell Correlation
resolution (1/A)

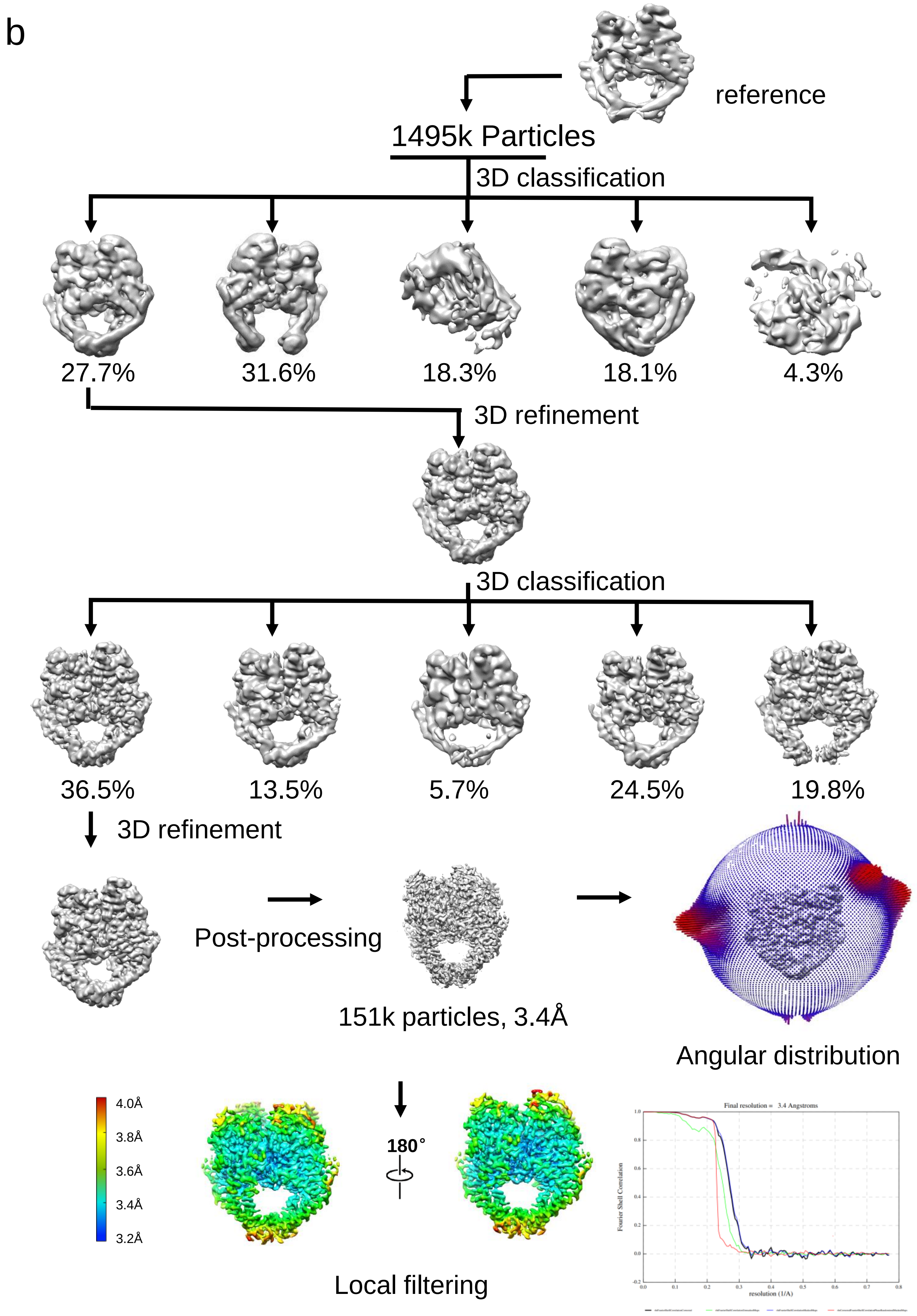
b
reference
1495k Particles
3D classification
27.7%
31.6%
18.3%
18.1%
4.3%
3D refinement
3D classification
36.5%
13.5%
5.7%
24.5%
19.8%
3D refinement
Post-processing
151k particles, 3.4Å
Angular distribution
4.0Å
3.8Å
3.6Å
3.4Å
3.2Å
180°
Local filtering
Final resolution = 3.4 Angstroms
Fourier Shell Correlation
resolution (1/A)

c

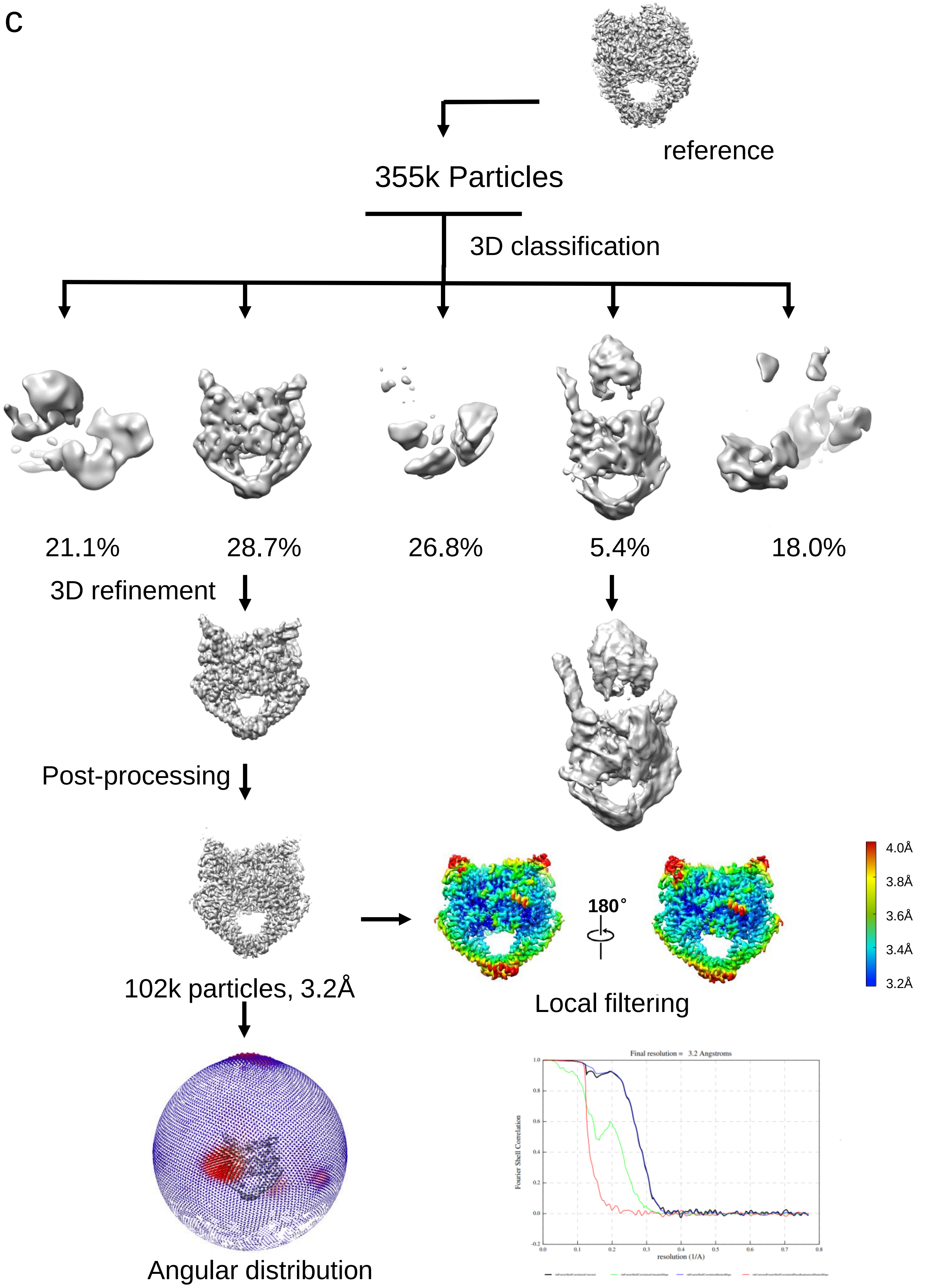
reference
355k Particles
3D classification
21.1%
28.7%
26.8%
5.4%
18.0%
3D refinement
Post-processing
102k particles, 3.2Å
180°
4.0Å
3.8Å
3.6Å
3.4Å
3.2Å
Local filtering
Angular distribution
Final resolution = 3.2 Angstroms
Fourier Shell Correlation
resolution (1/A)

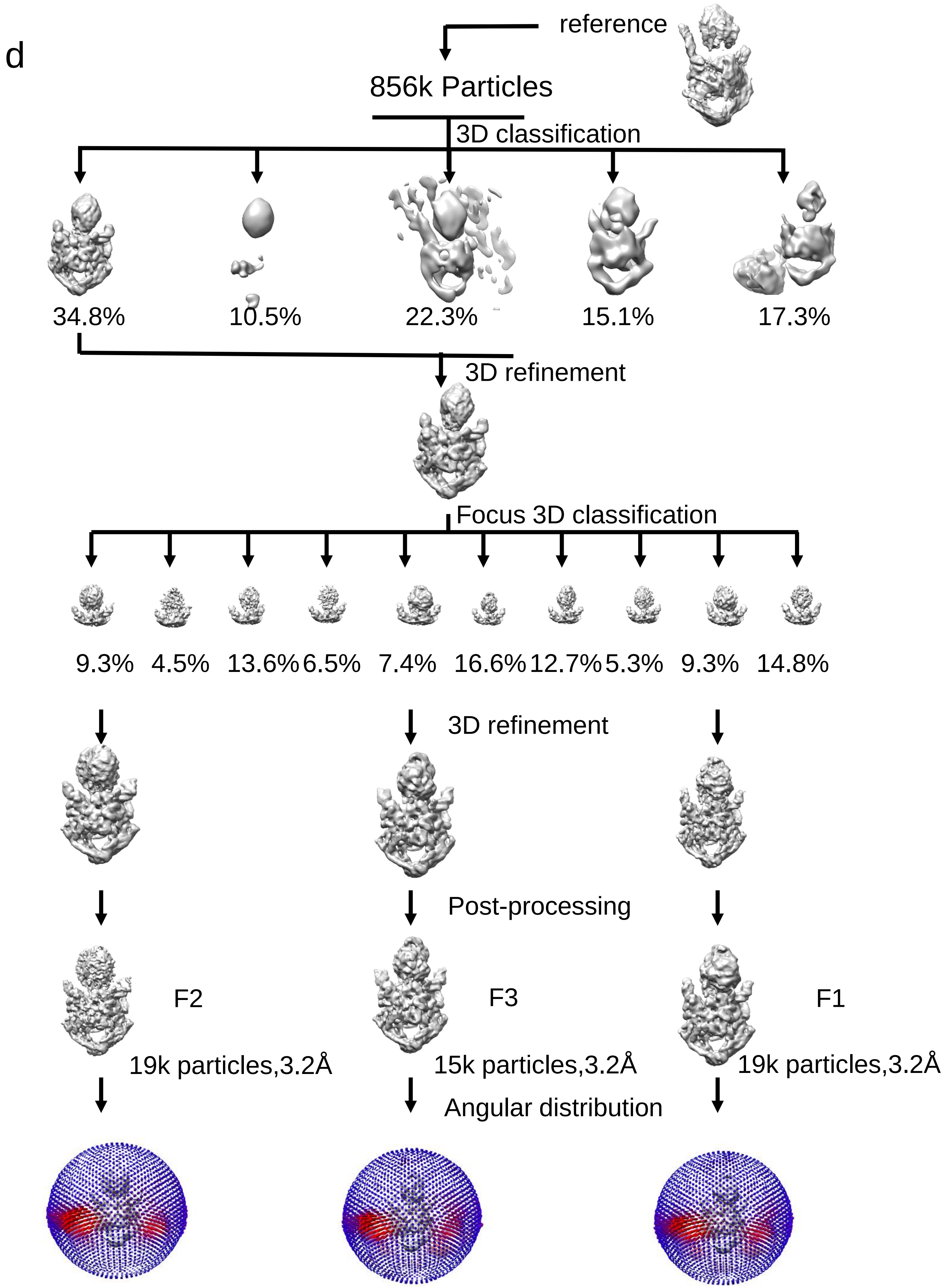
d
reference
856k Particles
3D classification
34.8%
10.5%
22.3%
15.1%
17.3%
3D refinement
Focus 3D classification
9.3%
4.5%
13.6%
6.5%
7.4%
16.6%
12.7%
5.3%
9.3%
14.8%
3D refinement
Post-processing
F2
F3
F1
19k particles,3.2Å
15k particles,3.2Å
19k particles,3.2Å
Angular distribution

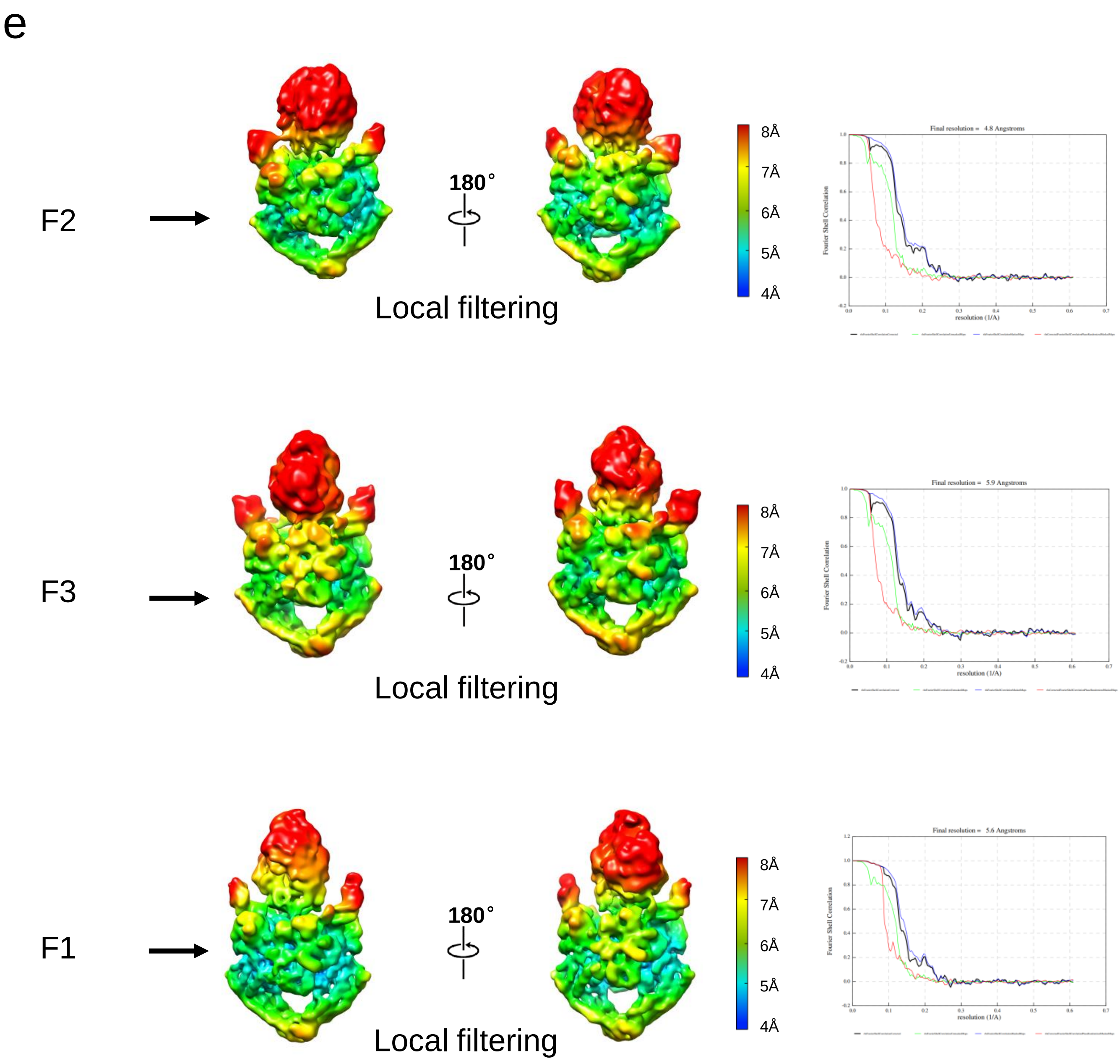

e
F2
180°
Local filtering
8Å
7Å
6Å
5Å
4Å
Final resolution = 4.8 Angstroms
Fourier Shell Correlation
resolution (1/A)
F3
180°
Local filtering
8Å
7Å
6Å
5Å
4Å
Final resolution = 5.9 Angstroms
Fourier Shell Correlation
resolution (1/A)
F1
180°
Local filtering
8Å
7Å
6Å
5Å
4Å
Final resolution = 5.6 Angstroms
Fourier Shell Correlation
resolution (1/A)

f

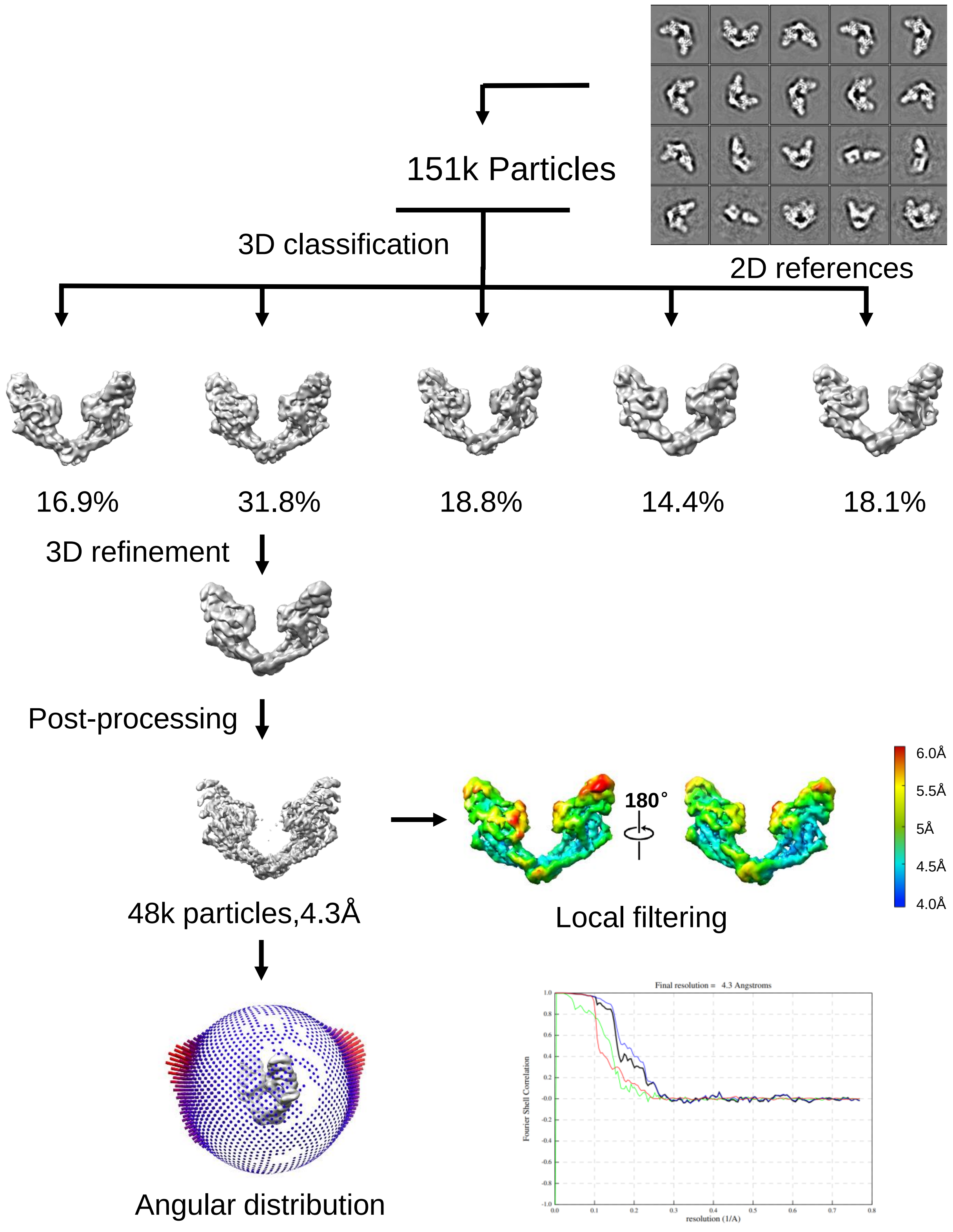
151k Particles
2D references
3D classification
16.9%
31.8%
18.8%
14.4%
18.1%
3D refinement
Post-processing
48k particles,4.3Å
180°
6.0Å
5.5Å
5Å
4.5Å
4.0Å
Local filtering
Angular distribution
Final resolution = 4.3 Angstroms
Fourier Shell Correlation
resolution (1/A)

f

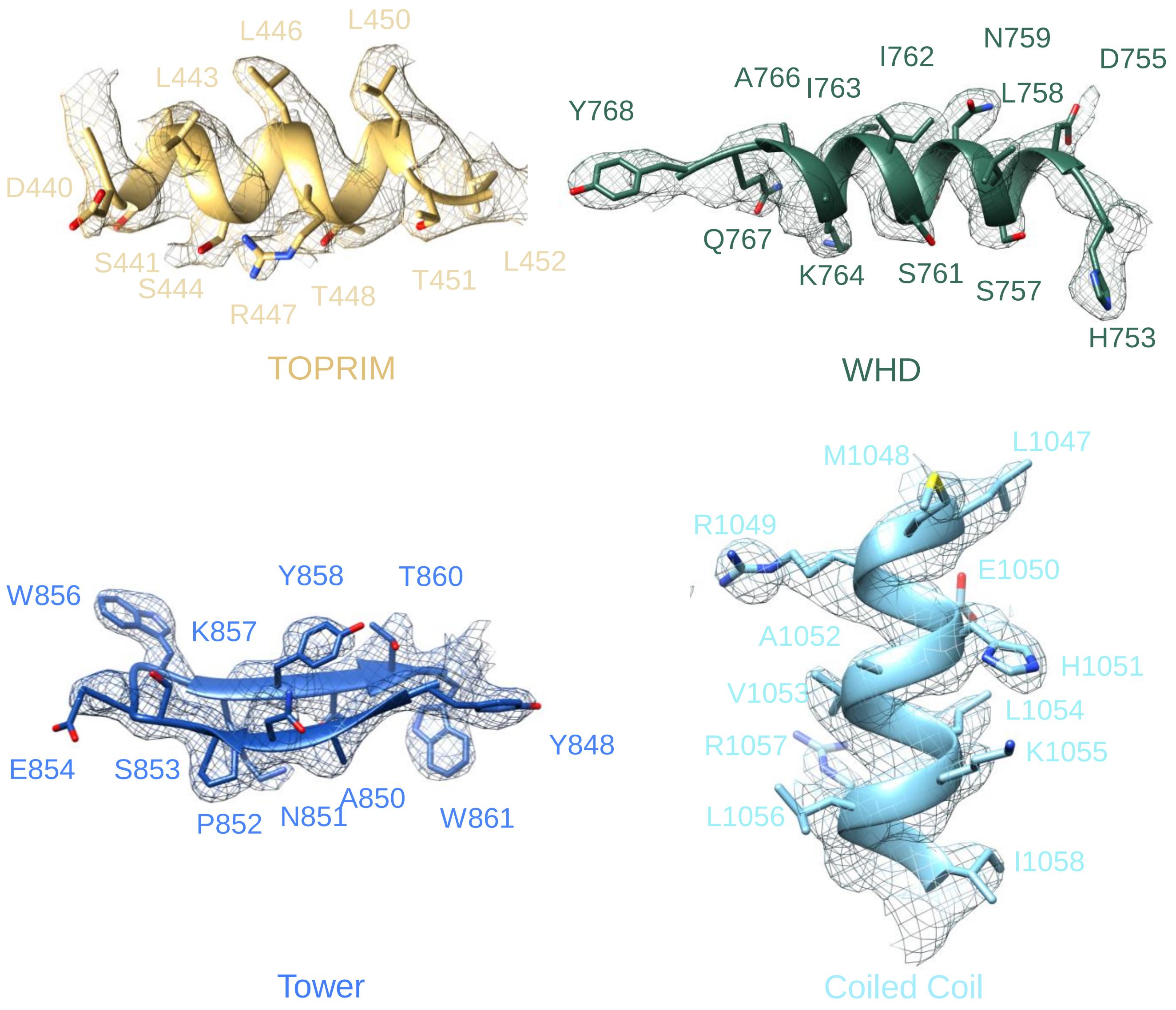

**Supplementary Figure 2: Cryo-EM analysis of the pP1192R and its complex. a-e** A schematic diagram illustrating the Cryo-EM data processing procedures for pP1192R and its complex. Local resolution, Euler angle distribution and the FSC curves for each reconstruction were demonstrated, respectively. **f** Representative localized density maps of pP1192R$_{CD\text{-}DNA}$ reconstruction.

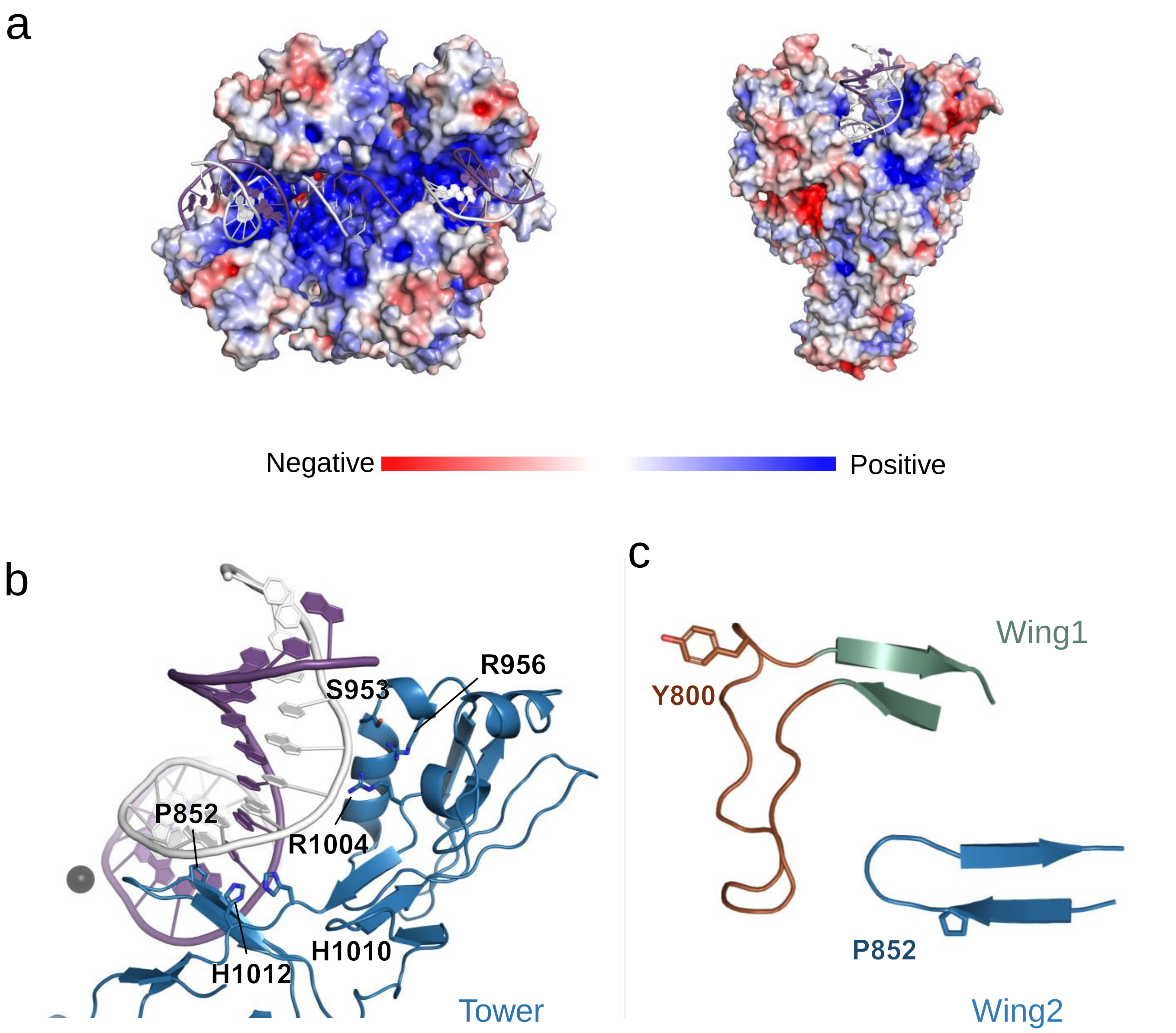

**Supplementary Figure 3: Interactions between G-segment DNA and pP1192R. a** Electrostatic potential map of the pP1192R$_{CD-DNA}$ structure from negative (red) to positive (blue), the DNA is shown as cartoon. **b** Magnified view of the interaction interfaces between the Tower subdomain and DNA. **c** Two conserved β-hairpins within the central domain of pP1192R.

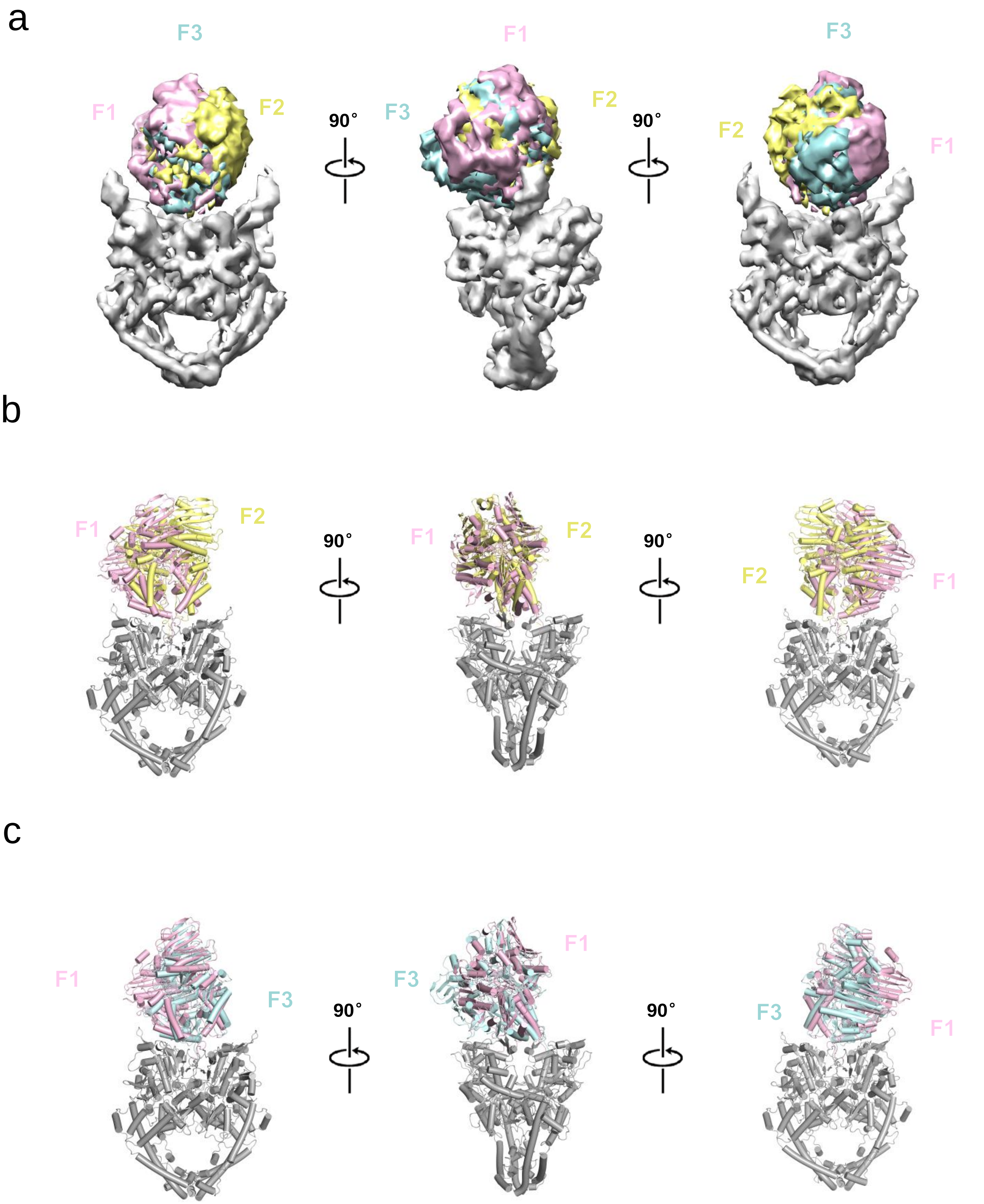
a
F3
F1
F2
90°
F1
F3
F2
90°
F3
F2
F1
b
F1
F2
90°
F1
F2
90°
F2
F1
c
F1
F3
90°
F1
F3
90°
F3
F1

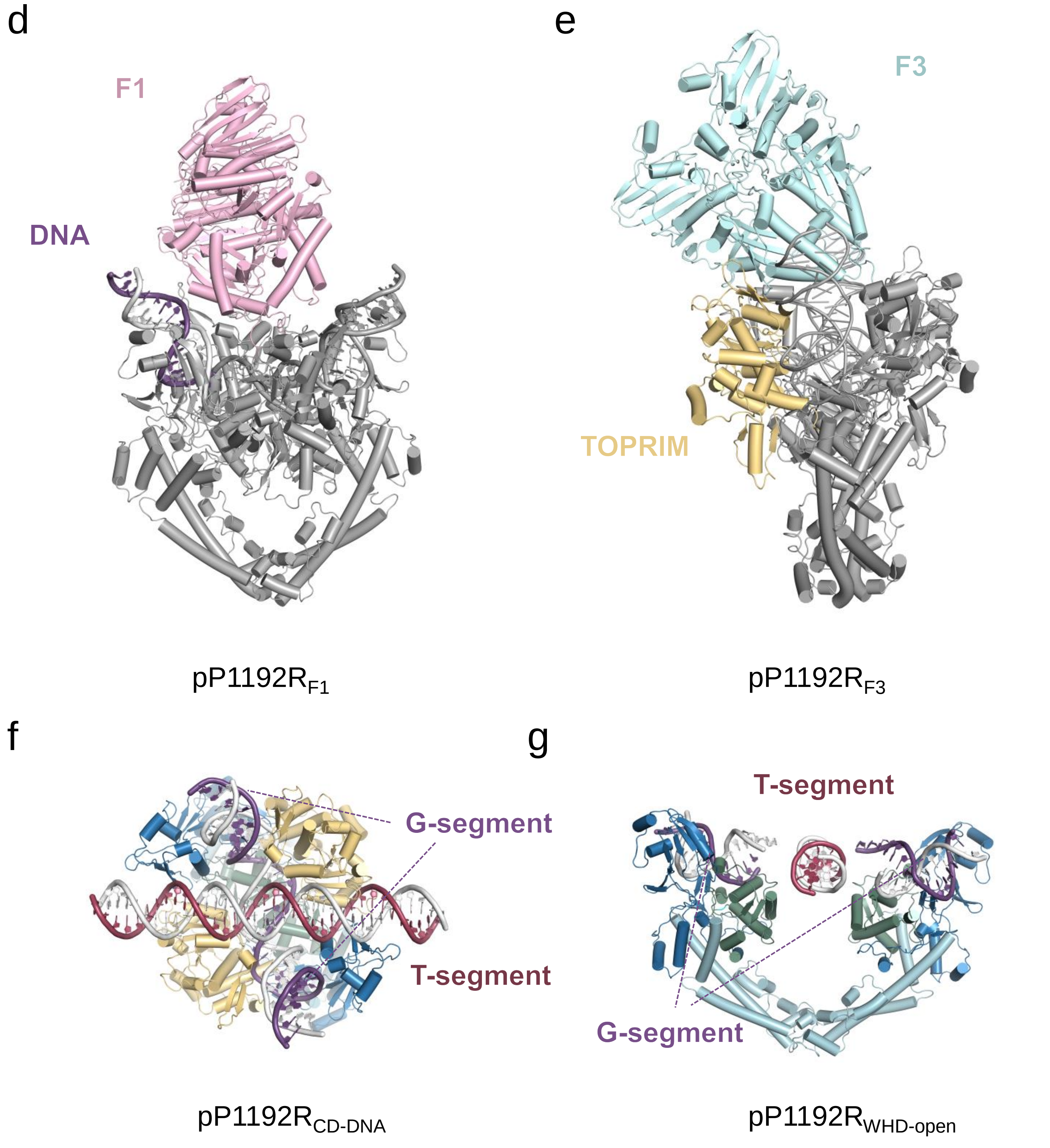

**Supplementary Figure 4: Conformational changes of pP1192R and pP1192R-DNA complex. a** Superimposition of $pP1192R_{F1-3}$ with distinct orientations of the ATPase domain. **b** Superimposition of $pP1192R_{F1}$ and $pP1192R_{F2}$ **c** Superimposition of $pP1192R_{F1}$ and $pP1192R_{F3}$. **d** The ATPase domain tilts towards the DNA terminal in $pP1192R_{F1}$ state. **e** The ATPase domain tilts towards the TOPRIM subdomain in $pP1192R_{F3}$ state, highlighted in yellow. **f** A modeled T-segment positioned above the G-segment within the $pP1192R_{CD-DNA}$ structure. **g** A modeled T-segment threading through the gap within the structure of $pP1192R_{WHD-open}$.

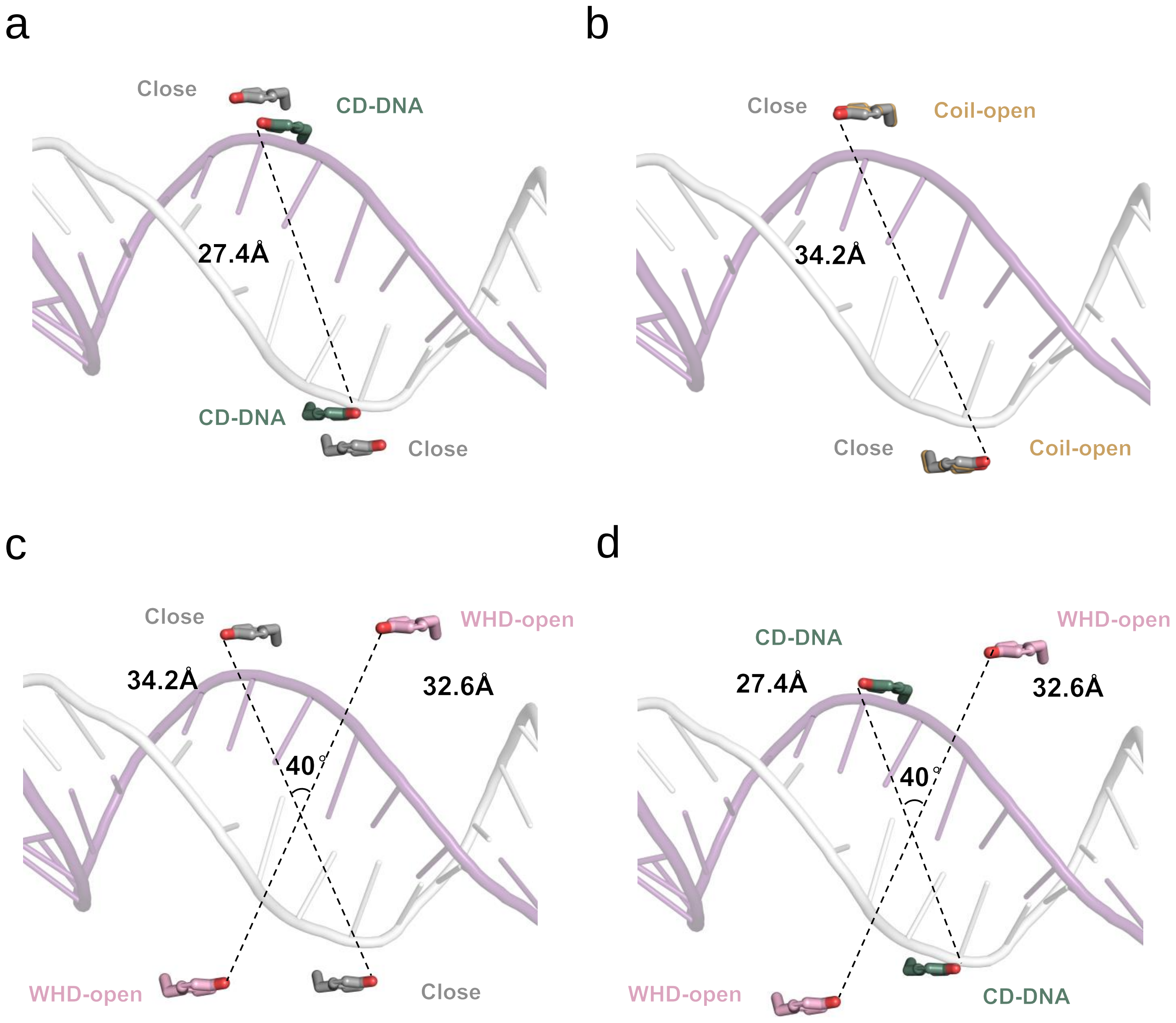

**Supplementary Figure 5: The positional variations between two catalytic Y800 in different conformations of pP1192R.** Comparison of the positional change between the two Y800 within pP1192R$_{CD\text{-}DNA}$ and pP1192R$_{Close}$ **(a)**, pP1192R$_{Coil\text{-}open}$ and pP1192R$_{Close}$ **(b)**, pP1192R$_{WHD\text{-}open}$ and pP1192R$_{Close}$ **(c)**, pP1192R$_{CD\text{-}DNA}$ and pP1192R$_{WHD\text{-}open}$ **(d)**, respectively.

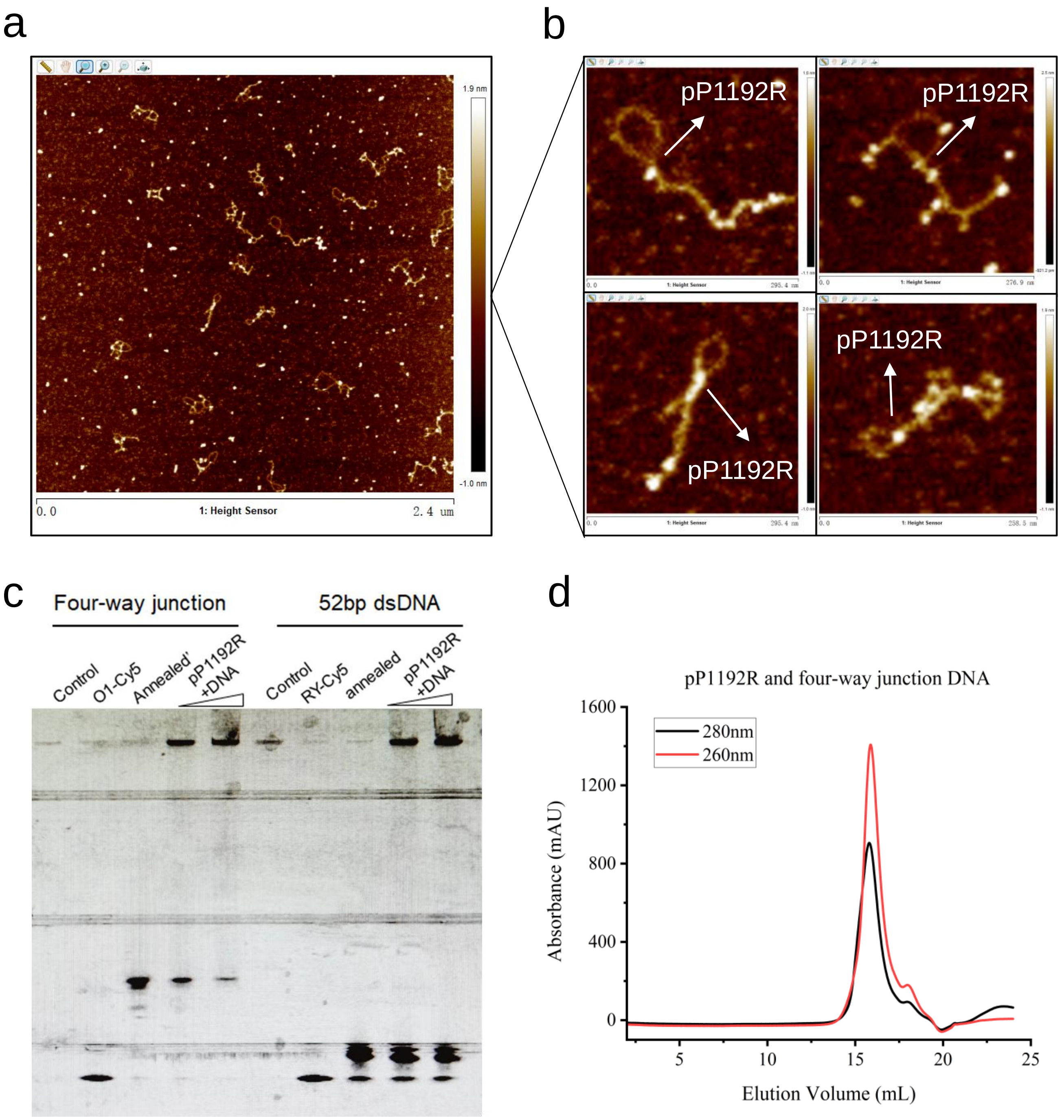

**Supplementary Figure 6: Interaction of pP1192R with DNA crossovers. a** AFM image of pP1192R and negatively supercoiled plasmid pUC19. **b** Close-up view of the interaction between pP1192R and pUC19. White arrows indicate the positions of pP1192R, which tend to bind to the DNA crossover rejoins. **c** Result of EMSA. Four-way junction and 52bp dsDNA are assayed respectively. Protein-bound DNA have a slow migration. "Control" represents pP1192R and "O1-Cy5'', "RY-Cy5" indicate 5'-labelled DNA Oligo controls. **d** Gel filtration of pP1192R and four-way junction DNA complex on a Superose 6 increase size-exclusion column. The ratio of absorbance values at 260nm and 280nm indicates a prominent nucleic acid binding.

```
              1        10                    20        30
ASFV          MEAFEISDFKEH.....................AKKKSMWAGALN.......KVTISGLM
S.cerevisiae  MSTEPVSASDK......................YQKISQLEHILKRPDTYIGSVETQEQL
H.sapiens     MEVSPLQPVNENMQVNKIKKNEDAKKRLSVERIYQKKTQLEHILLRPDTYIGSVELVTQQ

                   40        50        60        70        80        90
ASFV          GVFTEDEDLMALPIHRDHCPALLKIFDEIIVNATDHERACHNKTKKVTYIKISFDKGVFS
S.cerevisiae  QWIYDEETDCMIEKNVTIVPGLFKIFDEILVNAADNK..VRDPSMKRIDVNIHAEEHTIE
H.sapiens     MWVYDEDVG.INYREVTFVPGLYKIFDEILVNAADNK..QRDPKMSCIRVTIDPENNLIS

                   100       110       120       130       140       150
ASFV          CENDGPGIPIAKHEQASLIAKRDVYVPEVASCHFLAGTNINKAKDCIKGGTNGVGLKLAM
S.cerevisiae  VKNDGKGIPIEIHNKE......NIYIPEMIFGHLLTSSNYDDDEKKVTGGRNGYGAKLCN
H.sapiens     IWNNGKGIPVVEHKVE......KMYVPALIFGQLLTSSNYDDDEKKVTGGRNGYGAKLCN

                   160         170       180         190       200
ASFV          VHSQWAILTTADG..AQKYVQHINQRLDIIEPPTITPSR..EMFTRIELMPVYQELGYAE
S.cerevisiae  IFSTEFILETADLNVGQKYVQKWENNMSICHPPKITSYKKGPSYTKVTFKPDLTRFGMKE
H.sapiens     IFSTKFTVETASREYKKMFKQTWMDNMGRAGEMELKPFN.GEDYTCITFQPDLSKFKMQS

             210       220       230       240       250       260
ASFV          PLSETEQADLSAWIYLRACQCAAYVGKGTTIYYNDKPCRTGSVMALAKMYTLLSAPNSTI
S.cerevisiae  .....LDNDILGVMRRRVYDINGSV.RDINVYLNGKSLKIRNFKNYVELYLKSLEKKRQL
H.sapiens     .....LDKDIVALMVRRAYDIAGST.KDVKVFLNGNKLPVKGFRSYVDMYLK..DKLDET

             270            280       290       300       310       320
ASFV          HTAT.....IKADAKPYSLHPLQVAAVVSPKFKKFEHVSVINGVNCVKG.EHVTFLKKTI
S.cerevisiae  DNGEDGAAKSDIPTILYERINNRWEVAFAVSDISFQQISFVNSIATTMGGTHVNYITDQI
H.sapiens     GNS.........LKVIHEQVNHRWEVCLTMSEKGFQQISFVNSIATSKGGRHVDYVADQI

                   330       340       350       360       370       380
ASFV          NEMVVKKFQQTIKDKNRKTTLRDSCSNIFIVIVGSIPGIEWTGQRKDELSIAENVFKTHY
S.cerevisiae  VKKISEILKKKKKK..SVKSFQIKN.NMFIFINCLIENPAFTSQTKEQLTTRVKDFGSRC
H.sapiens     VTKLVDVVKKKNKGGVAVKAHQVKN.HMWIFVNALIENPTFDSQTKENMTLQPKSFGSTC

                   390       400                   410       420
ASFV          SIPSSFLTSMTKSIVDILLQSISKKD.............NHKQVDVDKYTRARNAGGKRA
S.cerevisiae  EIPLEYINKIMKTDLATRMFEIADANEENAL.KKSDGTRKSRITNYPKLEDANKAGTKEG
H.sapiens     QLSEKFIKAAIGCGIVESILNWVKFKAQVQLNKKCSAVKHNRIKGIPKLDDANDAGGRNS

            430       440       450       460       470       480
ASFV          QDCMLLAAEGDSALSLLRTGLTLGKSNPSGPSFDFCGMISLGGVIMNACKKVTNITTDSG
S.cerevisiae  YKCTLVLTEGDSALSLAVAGLAVVGR.......DYYGCYPLRGKMLNVREAS........
H.sapiens     TECTLILTEGDSAKTLAVSGLGVVGR.......DKYGVFPLRGKILNVREAS........

            490       500       510       520       530       540
ASFV          ETIMVRNEQLTNNKVLQGIVQVLGLDFNCHYKTQEERAKLRYGCIVACVDQDLDGCGKIL
S.cerevisiae  ......ADQILKNAEIQAIKKIMGLQHRKKYEDT...KSLRYGHLMIMTDQDHDGS.HIK
H.sapiens     ......HKQIMENAEINNIIKIVGLQYKKNYEDEDSLKTLRYGKIMIMTDQDQDGS.HIK

            550       560         570       580         590       600
ASFV          GLLLAYFHLFWPQLIIHG.FVKRLLTPLIRVYEKG.KTMPVEFYYEQEFDAWAKKQTSLA
S.cerevisiae  GLIINFLESSFPGLLDIQGFLLEFITPIIKVSITKPTKNTIAFYNMPDYEKWREEESHKF
H.sapiens     GLLINFIHHNWPSLLRHR.FLEEFITPIVKVSKNK...QEMAFYSLPEFEEWKSSTPNHK
```

```
              610       620       630       640       650       660
ASFV          NHTVKYYKGLAAHDTHEVKSMFKHFDNMVYTFTLDDS.AKELFHIYFGGESELRKRELCT
S.cerevisiae  TWKQKYYKGLGTSLAQEVREYFSNLDRHLKIFHSLQGNDKDYIDLAFSKKKADDRKEWLR
H.sapiens     KWKVKYYKGLGTSTSKEAKEYFADMKRHRIQFKYSGPEDDAAISLAFSKKQIDDRKEWLT

                670                 680       690       700       710
ASFV          GVVPLTET........QTQSIHSVRRIPCSLHLQVDTKAYKLDAIERQIPNFLDGMTRAR
S.cerevisiae  QYEPG............TVLDPTLKEIPISDFINKELILFSLADNIRSIPNVLDGFKPGQ
H.sapiens     NFMEDRRQRKLLGLPEDYLYGQTTTYLTYNDFINKELILFSNSDNERSIPSMVDGLKPGQ

            720       730       740       750       760       770
ASFV          RKILAGGVKCFASN.NRERKVFQFGGYVADHMFYHHGDMSLNTSIIKAAQYYPGSSHLYP
S.cerevisiae  RKVLYG...CFKKNLKSELKVAQLAPYVSECTAYHHGEQSLAQTIIGLAQNFVGSNNIY.
H.sapiens     RKVLFT...CFKRNDKREVKVAQLAGSVAEMSSYHHGEMSLMMTIINLAQNFVGSNNLN.

              780       790       800       810       820       830
ASFV          VFIGIGSFGSRHLGGKDAGSPRYISVQLASEFIKTMFPAEDSWLLPYVFEDGQRAEPEYY
S.cerevisiae  LLLPNGAFGTRATGGKDAAAARYIYTELN.KLTRKIFHPADDPLYKYIQEDEKTVEPEWY
H.sapiens     LLQPIGQFGTRLHGGKDSASPRYIFTMLS.SLARLLFPPKDDHTLKFLYDDNQRVEPEWY

              840       850       860       870       880       890
ASFV          VPVLPLAIMEYGANPSEGWKYTTWARQLEDILALVRAYVDKDNPKHELLHYAIKHKITIL
S.cerevisiae  LPILPMILVNGAEGIGTGWSTYIPPFNPLEIIKNIRHLMNDEEL...............E
H.sapiens     IPIIPMVLINGAEGIGTGWSCKIPNFDVREIVNNIRRLMDGEEP...............L

              900       910       920       930       940       950
ASFV          PLRPSNYNFKGHLKRFGQYYYSYGTYVISEQRNIITITELPLRVPTVAYIESIKKSSN..
S.cerevisiae  QMHPWFRGWTGTIEEIEPLRYRMYGRIEQIGDNVLEITELPARTWTSTIKEYLLLGLS.G
H.sapiens     PMLPSYKNFKGTIEELAPNQYVISGEVAILNSTTIEISELPVRTWTQTYKEQVLEPMLNG

                    960       970       980       990      1000      1010
ASFV          ...RMTFIEEIIDYSSSETIEILVKLKPNSLNRIVEEFKETEEQDSIENFLRLRNCLHSH
S.cerevisiae  NDKIKPWIKDMEEQHDDN.IKFIITLSP.......EEMAKTRKIGFYERFKLISPISLMN
H.sapiens     TEKTPPLITDYREYHTDTTVKFVVKMTE.......EKLAEAERVGLHKVFKLQTSLTCNS

                   1020      1030      1040      1050      1060      1070
ASFV          LNFVKPKGGIIEFNTYYEILYAWLPYRRELYQKRLMREHAVLKLRIIMETAIVRYINESA
S.cerevisiae  MVAFDPHGKIKKYNSVNEILSEFYYVRLEYYQKRKDHMSERLQWEVEKYSFQVKFIKMII
H.sapiens     MVLFDHVGCLKKYDTVLDILRDFFELRLKYYGLRKEWLLGMLGAESAKLNNQARFILEKI

                   1080      1090                          1100      1110
ASFV          ELNLSHYED.EKEASRILSEHGFP...............PLNHTLIISPEFASIEELN..
S.cerevisiae  EKELTVTNKPRNAIIQELENLGFPRFNKEGKPYYGSPNDEIAEQINDVKGATSDEEDEES
H.sapiens     DGKIIIENKPKKELIKVLIQRGYD.............SDPVKAWKEAQQKVPDEEENEES

                             1120      1130      1140      1150      1160
ASFV          ...........QKALQGCYTYILSLQARELLIAAKTRRVEKIKKMQARLDKVEQLLQESP
S.cerevisiae  SHEDTENVINGPEELYGTYEYLLGMRIWSLTKERYQKLLKQKQEKETELENLLKLSAKDI
H.sapiens     DNEKETEKSDSVTDSGPTFNYLLDMPLWYLTKEKKDELCRLRNEKEQELDTLKRKSPSDL

                   1170      1180      1190
ASFV          FPG.ASVWLEEIDAVEKAIIKGRNTQWKFH..............................
S.cerevisiae  WNTDLKAFEVGYQEFLQRDAEARGGNVPNKGSKTKGK............GKR...KLVDD
H.sapiens     WKEDLATFIEELEAVEAKEKQDEQVGLPGKGGKAKGKKTQMAEVLPSPRGQRVIPRITIE
```

```
ASFV          ............................................................
S.cerevisiae  EDYDPSKKNKKSTARKGKKIKLEDKNFERILLEQKLVTKSKAP..TKIKKEKTPSVSETK
H.sapiens     MKAEAEKKNKKKIKNENTEGSPQEDGVELEGLKQRLEKKQKREPGTKTKKQTTLAFKPIK

ASFV          ............................................................
S.cerevisiae  TEEEENAPSSTSSS....................................SIFDIKKEDK
H.sapiens     KGKKRNPWSDSESDRSSDESNFDVPPRETEPRRAATKTKFTMDLDSDEDFSDFDEKTDDE

ASFV          ............................................................
S.cerevisiae  D..........EGELSKISNK.FKKISTIFDKMGS.........TSATSKENTPEQDDVA
H.sapiens     DFVPSDASPPKTKTSPKLSNKELKPQKSVVSDLEADDVKGSVPLSSSPPATHFPDETEIT

ASFV          ............................................................
S.cerevisiae  T..KKNQTTAKKTAVKPKLAKKPVRKQQKVVELSGESDLEILDSYTDREDSNKDEDDAIP
H.sapiens     NPVPKKNVTVKKTAAKSQSSTSTTGAKKRAAPKGTKRDPALNSGVSQKPDPAKTKNRRKR

ASFV          ............................................................
S.cerevisiae  QRSRRQRS........SRAASVPK.KSYVETLELSDDSFIEDDEEENQGSD.VSFNEED.
H.sapiens     KPSTSDDSDSNFEKIVSKAVTSKKSKGESDDFHMDFDSAVAPRAKSVRAKKPIKYLEESD

ASFV          .....
S.cerevisiae  .....
H.sapiens     EDDLF
```

**Supplementary Figure 7: Sequence alignment of pP1192R with Topo II of *S. cerevisiae* and *H. sapiens* .**

**Supplementary Table 1: Statistics of Data Collection, Image Processing and Model Building.**

| | CD-DNA | F1 | F2 | F3 |
|---|---|---|---|---|
| **Data collection** | | | | |
| **EM equipment** | FEI Titan Krios | FEI Titan Krios | FEI Titan Krios | FEI Titan Krios |
| **Voltage (kV)** | 300 | 300 | 300 | 300 |
| **Detector** | Gatan K2 | Gatan K2 | Gatan K2 | Gatan K2 |
| **Pixel size (Å/pixel)** | 0.65 | 0.82 | 0.82 | 0.82 |
| **Electron dose ($e^-$/Å2)** | 60 | 60 | 60 | 60 |
| **Defocus range (μm)** | -1.2-2.5 | -1.2-2.5 | -1.2-2.5 | -1.2-2.5 |
| **Reconstruction** | | | | |
| **Software** | Relion3.0 | Relion3.0 | Relion3.0 | Relion3.0 |
| **Number of used particles** | 102k | 19k | 19k | 15k |
| **Symmetry** | C1 | C1 | C1 | C1 |
| **Map sharpening B-factor ($Å^2$)** | -106 | -80 | -80 | -80 |
| **Final resolution (Å)** | 3.2 | 5.6 | 4.8 | 5.9 |
| **Model building** | | | | |
| **Software** | Coot | Coot | Coot | Coot |
| **Model Refinement** | | | | |
| **Software** | PHENIX | PHENIX | PHENIX | PHENIX |
| **Map CC (mask)** | 0.836 | 0.608 | 0.459 | 0.578 |
| **Map CC (peaks)** | 0.712 | 0.517 | 0.415 | 0.482 |
| **Map CC (volume)** | 0.811 | 0.597 | 0.482 | 0.562 |
| **Rmsd (bonds) (Å)** | 0.0084 | 0.0088 | 0.0088 | 0.0088 |
| **Rmsd (angles) (°)** | 1.21 | 1.24 | 1.24 | 1.25 |
| **Validation** | | | | |
| **MolProbity score** | 1.49 | 1.99 | 1.84 | 1.86 |
| **Clash score** | 3.91 | 13.87 | 11.80 | 11.79 |
| **Ramachandran plot** | | | | |
| **Outliers (%)** | 0.0 | 0.0 | 0.0 | 0.0 |
| **Allowed (%)** | 4.1 | 3.5 | 3.2 | 3.2 |
| **Favored (%)** | 95.9 | 96.5 | 96.8 | 96.8 |
| **Rotamer outliers (%)** | 0.15 | 1.15 | 0.73 | 0.15 |
| **Cβ outliers (%)** | 0 | 0 | 0 | 0 |

**Supplementary Table 1: Statistics of Data Collection, Image Processing and Model Building.**

| | Coil-open | Close | WHD-open |
|---|---|---|---|
| **Data collection** | | | |
| **EM equipment** | FEI Titan Krios | FEI Titan Krios | FEI Titan Krios |
| **Voltage (kV)** | 300 | 300 | 300 |
| **Detector** | Gatan K2 | Gatan K2 | Gatan K2 |
| **Pixel size (Å/pixel)** | 0.65 | 0.65 | 0.65 |
| **Electron dose ($e^-$/Å2)** | 60 | 60 | 60 |
| **Defocus range (μm)** | -1.2-2.5 | -1.2-2.5 | -1.2-2.5 |
| **Reconstruction** | | | |
| **Software** | Relion3.0 | Relion3.0 | Relion3.0 |
| **Number of used particles** | 150k | 102k | 48k |
| **Symmetry** | C2 | C2 | C2 |
| **Map sharpening B-factor (Å$^2$)** | -148 | -160 | -201 |
| **Final resolution (Å)** | 3.3 | 3.4 | 4.3 |
| **Model building** | | | |
| **Software** | Coot | Coot | Coot |
| **Model Refinement** | | | |
| **Software** | PHENIX | PHENIX | PHENIX |
| **Map CC (mask)** | 0.848 | 0.825 | 0.769 |
| **Map CC (peaks)** | 0.689 | 0.654 | 0.679 |
| **Map CC (volume)** | 0.824 | 0.789 | 0.758 |
| **Rmsd (bonds) (Å)** | 0.0110 | 0.0109 | 0.0076 |
| **Rmsd (angles) (°)** | 1.31 | 1.40 | 1.25 |
| **Validation** | | | |
| **MolProbity score** | 1.66 | 1.53 | 1.92 |
| **Clash score** | 4.97 | 2.80 | 8.04 |
| **Ramachandran plot** | | | |
| **Outliers (%)** | 0.0 | 0.0 | 0.0 |
| **Allowed (%)** | 4.9 | 6.1 | 5.6 |
| **Favored (%)** | 95.1 | 93.9 | 94.4 |
| **Rotamer outliers (%)** | 0.76 | 0.54 | 0.00 |
| **Cβ outliers (%)** | 0 | 0 | 0 |

**Supplementary Table 1: Statistics of Data Collection, Image Processing and Model Building.**

| | ATPase-AMPPNP | ATPase-ADP |
|---|---|---|
| **Data collection** | | |
| **Space group** | $P6_122$ | $P6_122$ |
| **Cell dimensions** | | |
| **a, b, c (Å)** | 85.08, 85.08, 210.18 | 85.23, 85.23, 209.81 |
| **α, β, γ (°)** | 90, 90, 120 | 90, 90, 120 |
| **Resolution (Å)** | 50.00-2.60 (2.64-2.60)[a] | 50.00-2.30 (2.34-2.30) |
| **$R_{merge}$** | 0.203 (1.019) | 0.149 (0.945) |
| **I/σI** | 11.00 (1.38) | 14.22 (2.24) |
| **Completeness (%)** | 100.0 (99.3) | 99.4 (99.4) |
| **Redundancy** | 19.0 (9.0) | 8.2 (7.7) |
| **No. of reflections** | 277,001 | 170,228 |
| **No. of unique reflections** | 14,604 | 20,799 |
| **Refinement** | | |
| **$R_{work}$/ $R_{free}$** | 0.200/0.244 | 0.193/0.246 |
| **No. of atoms** | | |
| **Protein** | 3,141 | 3,080 |
| **Ligand/ion** | 32 | 27 |
| **Water** | 121 | 108 |
| **Average B-factors** | | |
| **Protein** | 37.0 | 43.7 |
| **Ligand/ion** | 24.7 | 34.1 |
| **Water** | 31.9 | 38.1 |
| **R.m.s deviations** | | |
| **Bond lengths (Å)** | 0.010 | 0.004 |
| **Bond angles (°)** | 1.060 | 1.135 |
| **Ramachandran plot (%)** | | |
| **Favoured** | 98.5 | 98.5 |
| **Allowed** | 1.5 | 1.2 |
| **Outliers** | 0 | 0.3 |

[a]Values in parentheses correspond to the shell of the highest resolution

**Supplementary Table 3: Summary of the models**

| Subunit Name | Chain | Total residues/ range built | Unmodelled residues | % atomic model |
|---|---|---|---|---|
| **pP1192R$_{CD\text{-}DNA}$** | | | | |
| pP1192R | A | 1192/413-1192 | 1-412 | 780/1192 |
| pP1192R | B | 1192/415-1192 | 1-414 | 778/1192 |
| DNA | C | 52/17-48 | 1-16, 49-52 | 32/52 |
| DNA | D | 52/5-37 | 1-4, 38-52 | 33/52 |
| **pP1192R$_{F1}$** | | | | |
| pP1192R | A | 1192/3-1192 | 1-2 | 1190/1192 |
| pP1192R | B | 1192/3-1192 | 1-2 | 1190/1192 |
| DNA | C | 52/14-51 | 1-13, 52 | 38/52 |
| DNA | D | 52/4-41 | 1-3, 42-52 | 38/52 |
| **pP1192R$_{F2}$** | | | | |
| pP1192R | A | 1192/3-403, 413-1192 | 1-2, 404-412 | 1181/1192 |
| pP1192R | B | 1192/3-410, 415-1192 | 1-2, 411-414 | 1186/1192 |
| DNA | C | 52/14-51 | 1-13, 52 | 38/52 |
| DNA | D | 52/4-41 | 1-3, 42-52 | 38/52 |
| **pP1192R$_{F3}$** | | | | |
| pP1192R | A | 1192/3-1192 | 1-2 | 1190/1192 |
| pP1192R | B | 1192/3-405, 416-1192 | 1-2, 406-415 | 1180/1192 |
| DNA | C | 52/14-51 | 1-13, 52 | 38/52 |
| DNA | D | 52/4-41 | 1-3, 42-52 | 38/52 |

### Supplementary Table 3: Summary of the models

| Subunit Name | Chain | Total residues/ range built | Unmodelled residues | % atomic model |
|---|---|---|---|---|
| | | **pP1192R$_{\text{Coil-open}}$** | | |
| pP1192R | A | 1192/415-472, 501-893, 895-1192 | 1-414, 473-500, 894 | 749/1192 |
| pP1192R | B | 1192/415-472, 501-893, 895-1192 | 1-414, 473-500, 894 | 749/1192 |
| | | **pP1192R$_{\text{Close}}$** | | |
| pP1192R | A | 1192/415-473, 502-852, 856-893, 895-1192 | 1-414, 474-501, 853-855, 894 | 746/1192 |
| pP1192R | B | 1192/415-473, 502-852, 856-893, 895-1192 | 1-414, 474-501, 853-855, 894 | 746/1192 |
| | | **pP1192R$_{\text{WHD-open}}$** | | |
| pP1192R | A | 1192/701-1192 | 1-700 | 492/1192 |
| pP1192R | B | 1192/701-1192 | 1-700 | /1492192 |
| | | **ATPase-AMPPNP** | | |
| pP1192R | A | 434/3-405 | 1-2, 406-434 | 403/434 |
| | | **ATPase-ADP** | | |
| pP1192R | A | 434/3-334, 342-405 | 1-2, 335-341 | 396/434 |

### Supplementary Table 4: DNA oligonucleotides

| **52bp dsDNA** |
| --- |
| 5’-ATGCATATATATGTATATGTATGTGTGTATATAT ACACATATATATATATATAT-3’ |
| 5’-ATATATATATATATATGTGTATATATACACACATAC ATATACATATATATGCAT-3’ |

| **Four-way junction DNA** |
| --- |
| O1: 5’-CTGGACGCAATCTGACAATGCGCTCATCGTC ATCCTCGGCACGCGCCG-3’ |
| O2: 5’-CGGCGCGTGCCGAGGATGACGATGAGATAG GCGTTAACGCGGCCTA-3’ |
| O3: 5’-TAGGCCGCGTTAACGCCTATTTGCCCGGGAG TACCGGCATTCCT-3’ |
| O4: 5’-AGGAATGCCGGTACTCCCGGGCAACGCATTG TCAGATTGCGTCCAG-3’ |
